# Bromodomain Protein, BRD4, Contributes to the Regulation of Alternative Splicing

**DOI:** 10.1101/440297

**Authors:** Sheetal Uppal, Qingrong Chen, Daoud Meerzaman, Anne Gegonne, Dinah S. Singer

**Author notes:** Current address: Homi Bhabha National Institute (HBNI), Bhabha Atomic Research Centre (BARC), Trombay, Mumbai-400085,_India. ***Classification:*** Biological Sciences; biochemistry.

## Abstract

Bromodomain protein 4 (BRD4) is an atypical kinase and a histone acetyl transferase (HAT) which plays an important role in chromatin remodeling and early transcriptional elongation. During transcription elongation, BRD4 travels with the elongation complex. Since most of the alternative splicing events take place co-transcriptionally, we asked if BRD4 plays a role in regulation of alternative splicing. We find that distinct patterns of alternative splicing are associated with conditional deletion of BRD4 during thymocyte differentiation in vivo. Similarly, depletion of BRD4 in T-ALL cells alters patterns of splicing. Most of the alternatively spliced events affected by BRD4 are usage of exon skipping. In an established insulin receptor minigene model of splicing, BRD4 over expression modulates alternative splicing. Importantly, as assessed by both immunoprecipitation (IP) and proximity ligation (PLA) assays, BRD4 interacts with components of the splicing machinery. BRD4 also co-localizes on chromatin with one of the splicing regulators. We propose that BRD4 contributes to patterns of alternative splicing through its interaction with the splicing machinery during transcription elongation.

**Significance Statement:** The bromodomain protein, BRD4, is a transcriptional and epigenetic regulator that plays a critical role in both cancer and inflammation. It has pleiotropic activities, including chromatin organization, transcriptional pause release and initiation. We now report that it also contributes to the regulation of alternative splicing. Taken together, these findings indicate that BRD4 functions to coordinate the various steps in gene expression.

## Introduction

The bromodomain protein, BRD4, is a transcriptional and epigenetic regulator that plays a critical role in both cancer and inflammation (1). In addition to functioning as a passive scaffold that recruits a number of different transcription factors to promoters, BRD4 possesses both histone acetyl transferase (HAT) enzymatic activity and atypical kinase activity (2, 3). These enzymatic activities allow BRD4 to uniquely coordinate chromatin organization and transcription. Transcriptional activation requires chromatin remodeling through acetylation of nucleosomal histones and chromatin de-compaction around targeted genes. BRD4 HAT activity acetylates both histone tails and the globular domain of histone H3, resulting in nucleosome dissociation and chromatin remodeling at BRD4-target genes (2). BRD4’s intrinsic kinase activity phosphorylates Ser2 of the Pol II carboxy terminal domain (CTD) which hyperactivates Topoisomerase I, releasing the torsional stress imposed by early transcription elongation and facilitating pause release (3, 4). Thus, BRD4 coordinates chromatin organization and transcription by a stepwise process regulated through its pleiotropic enzymatic activities.

Importantly, phosphorylation of Ser2 of the CTD functions as a docking site for splicing factors (5). The majority of splicing in mammalian systems is mediated by the U2-dependent spliceosome, a multi-megadalton complex that assembles on intronic 5’ and 3’ splice sites (6). Components of the spliceosome include the U1-U6 RNPs. The activity of the spliceosome is regulated by a series of trans-acting factors, such as the HnRNP and SR protein families, that function to regulate alternative splicing. Although it was originally thought that alternative splicing largely occurred post-transcriptionally, it is now clear that most alternative splicing events are co-transcriptional (7, 8). Indeed, the two processes are closely linked and reciprocally regulate each other (9). RNA polymerase II (Pol II) plays a critical role in regulating alternative splicing, both in recruiting splicing factors and in defining exons. A number of regulators of alternative splicing, such as Fus, bind to the Ser2 phosphorylated form of the Pol II CTD (10). Elongation of Pol II along the body of the gene contributes to the regulation of alternative splicing (5). Pausing of Pol II in exons at the 3’ splice site allows recognition and processing by the splicing machinery. Splicing is also kinetically regulated by the rate of Pol II elongation, such that changes in the rate of elongation alter the patterns of splicing (5).

Factors that regulate transcription, such as chromatin remodelers and transcription factors further modulate splicing. Various transcription factors including Mediator, YB1, CTCF and Pu.1 have also been implicated in alternative splicing (11–14). Among the chromatin remodelers that contribute to alternative splicing patterns are the histone chaperone FACT and the ATP-dependent SWI/SNF complex (15, 16). Consistent with chromatin remodelers regulating splicing, nucleosome occupancy contributes to the determination of splicing patterns: Nucleosome density is greater on exons than on introns, thereby defining potential sites of splicing (17). Finally, histone acetylation and methylation patterns affect splicing patterns (18, 19).

Given the intimate relationship between splicing, transcription and chromatin structure and the dual functions of many transcription factors in both processes, we considered the possibility that BRD4 contributes to the regulation of alternative splicing. Lending support to this hypothesis was the finding that the BET protein, BRDT, forms a complex with spliceosome components (20). Furthermore, a large number of splicing factors, including Fus, U2AF2 and members of the HnRNP and SR families co-immunoprecipitate with BRD4 (21) and BRD4 depletion alters splicing patterns in response to heat shock (22).

Here we report that BRD4 modulates alternative splicing. *In vivo* deletion of BRD4 during thymocyte differentiation, affected the patterns of alternative splicing in each of the developmental thymocyte subpopulations. Similarly, depletion of BRD4 in T-ALL cells altered patterns of splicing. Using an insulin receptor mini-gene reporter system, co-transfection of BRD4 in HeLa cells led to an altered pattern of mini-gene transcript splicing. Significantly, BRD4 was shown to interact directly with components of the splicing machinery *in vitro* and to co-localize with them *in vivo.* Taken together, these findings demonstrate that BRD4 contributes to establishing patterns of alternative splicing.

## Results

### BRD4 deficiency in thymocytes alters alternative splicing patterns

The finding that BRD4 travels with the transcription elongation complex (23, 24) raised the question of its role during splicing. While BRD4 HAT activity reduces nucleosome occupancy across active genes and its kinase activity promotes transcription (2, 3), we considered the possibility that it is also involved, directly or indirectly, in RNA splicing. To determine whether BRD4 affects splicing patterns of endogenous genes *in vivo,* we examined the effect of deleting BRD4 on splicing during thymopoiesis. Our recent studies have identified a unique role for BRD4 in regulating patterns of gene expression during thymocyte differentiation (25). The generation of T cells in the thymus results from the sequential differentiation of a series of thymocyte precursors (Fig. 1A). The earliest thymic immigrants from the bone marrow do not express the markers associated with mature T cells, namely the CD4/CD8 coreceptor molecules, and are called Double Negatives (DN). DN thymocytes differentiate into CD4+CD8+ double positive (DP) thymocytes via an immature transitional cell that expresses only CD8 (immature single positive, ISP). DP thymocytes differentiate into the mature CD4+ and CD8+ single positive thymocytes that seed the periphery. BRD4 is expressed in all five subpopulations, at approximately equal levels. Deletion of BRD4 at the DN stage leads to a selective defect in maturation at the ISP stage (25).

**Figure 1.**
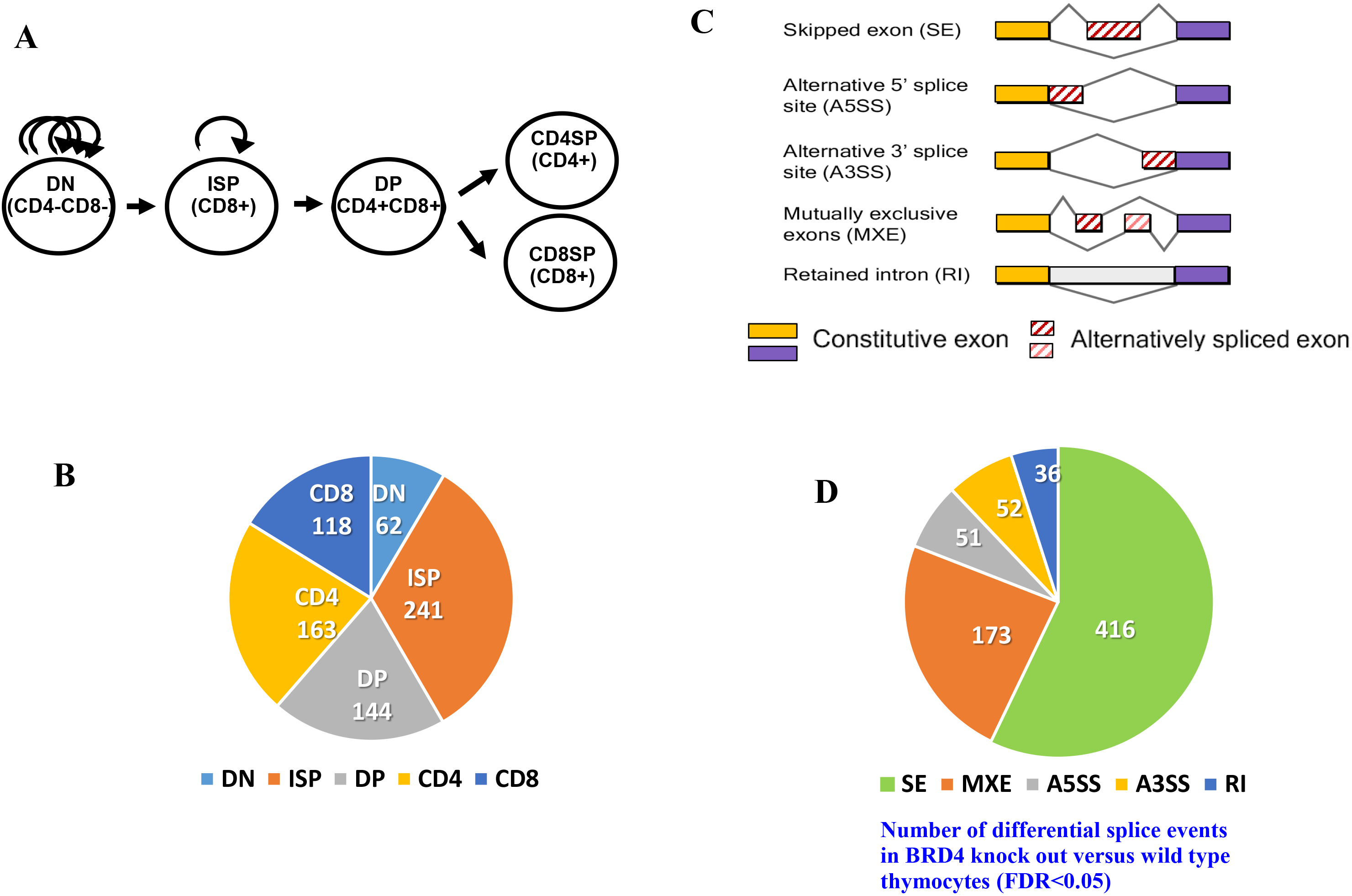
BRD4 regulates alternative splicing in murine thymocytes. (A) Developmental stages in thymocyte differentiation, DN (CD4, CD8 double negative), ISP (CD8+ immature single positive), DP (CD4, CD8 double positive), CD4 and CD8 single positive thymocytes are shown. Arched arrows denote level of proliferative activity in DN and ISP thymocytes. (B) Pie chart showing the number of differential splice events in different thymocytes subpopulations in BRD4 knock out versus wild type thymus (FDR<0.05), derived from RNA-seq analysis. BRD4 was conditionally deleted in DN thymocytes by LCK-Cre (25). (C) Schematic diagram depicting different types of alternatively spliced events (D) Pie chart showing the distribution of different types of alternatively spliced events among all of the differentially spliced events in total thymocytes from BRD4 knock out versus wild type (FDR<0.05).

To assess the effect of BRD4 on alternative splicing, we analyzed RNA-seq data from either wild-type or BRD4-deficient thymocytes purified at each of these developmental stages (25). As summarized in Fig. 1B, BRD4 deficiency affected alternative splicing patterns in each of the subpopulations. Notably, within the ISP population, there were 241 differences in splicing events between the BRD4-deficient ISP and wild type; splicing event differences also occurred in the other subpopulations as a result of BRD4 deletion. Consistent with BRD4 selectively affecting maturation of the ISP subpopulation (25), the largest differences occurred in that population.

Five different forms of alternative splicing have been defined: skipped exon (SE), alternative 5’ (A5SS) or alternative 3’ (A3SS) splice sites, mutually exclusive exons (MXE) and retained intron (RI) (Fig. 1C). Of these forms, the skipped exon was the predominant form affected by BRD4 deletion in thymocytes, although all of the forms were affected to some extent (Fig. 1D, Table 1). These data demonstrate that BRD4 contributes to splicing of endogenous genes *in vivo.*

**Table 1.**
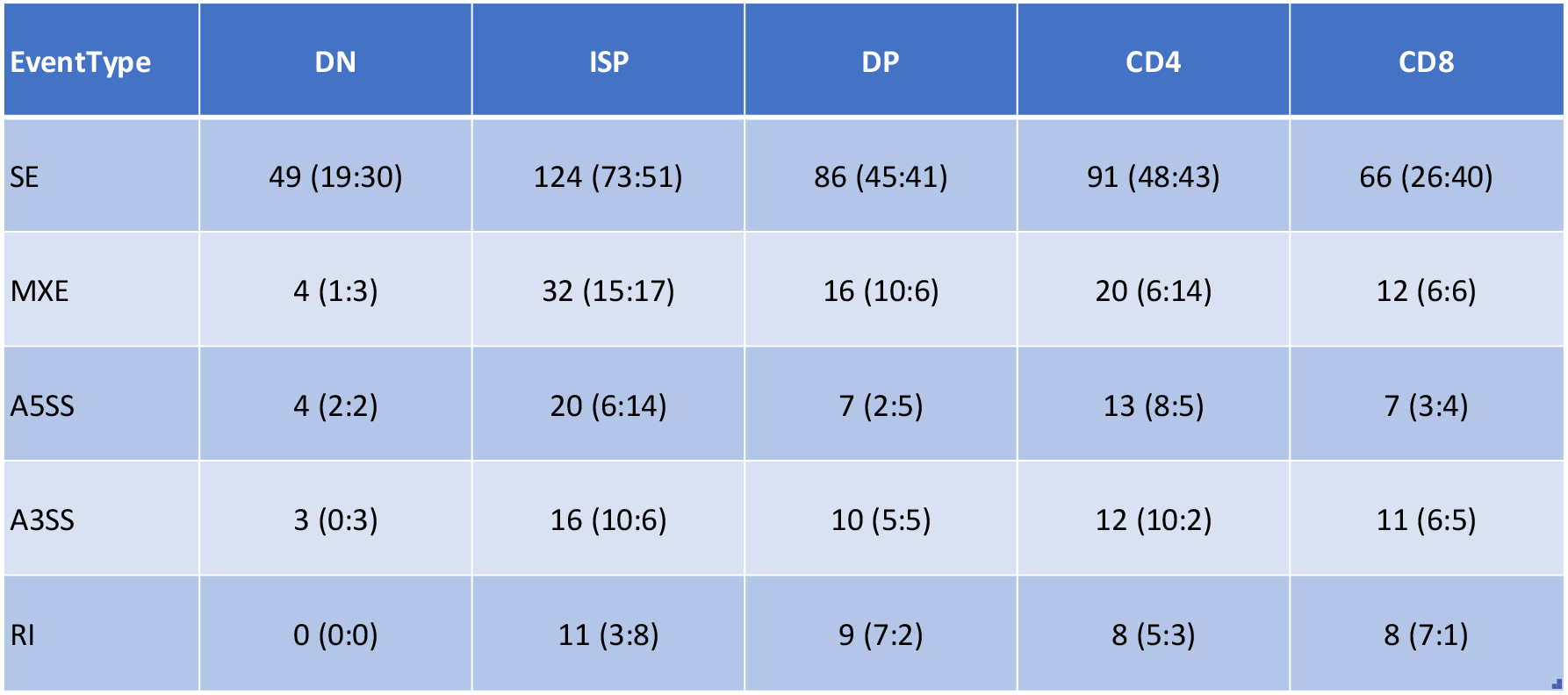
Deletion of BRD4 Results in Differential Splicing in Thymocyte Subpopulations Relative to WT mice. **Legend to Table 1** Shown are the numbers of significant events detected using Junction Counts (FDR<0.05 & |IncLevelDifference|>0.1). The numbers in the parentheses (n1:n2) represent the significant number of up-regulated events (high in KO) and down-regulated events (high in WT).

| EventType | DN | ISP | DP | CD4 | CD8 |
| --- | --- | --- | --- | --- | --- |
| SE | 49 (19:30) | 124 (73:51) | 86 (45:41) | 91 (48:43) | 66 (26:40) |
| MXE | 4 (1:3) | 32 (15:17) | 16 (10:6) | 20 (6:14) | 12 (6:6) |
| A5SS | 4 (2:2) | 20 (6:14) | 7 (2:5) | 13 (8:5) | 7 (3:4) |
| A3SS | 3 (0:3) | 16 (10:6) | 10 (5:5) | 12 (10:2) | 11 (6:5) |
| RI | 0 (0:0) | 11 (3:8) | 9 (7:2) | 8 (5:3) | 8 (7:1) |

Experimental analysis of the splicing patterns of individual genes within the thymocyte subpopulations validated the differences in splicing derived from the RNA seq data among the wild type subpopulations, as well as between wild type and BRD4-deficient cells in the same subpopulation (Fig. 2). In wild type thymocytes, exon 5 of CD45 is preferentially included in DN and ISP thymocytes but preferentially excluded in DP and CD4+ thymocytes (Fig. 2A,B). Similarly, exon 15 of Arhgef1 is preferentially excluded in all WT subpopulations except in CD8+ thymocytes. Exon 13 of Picalm is preferentially included in all wild type subpopulations except DP (Fig. 2C-F).

**Figure 2.**
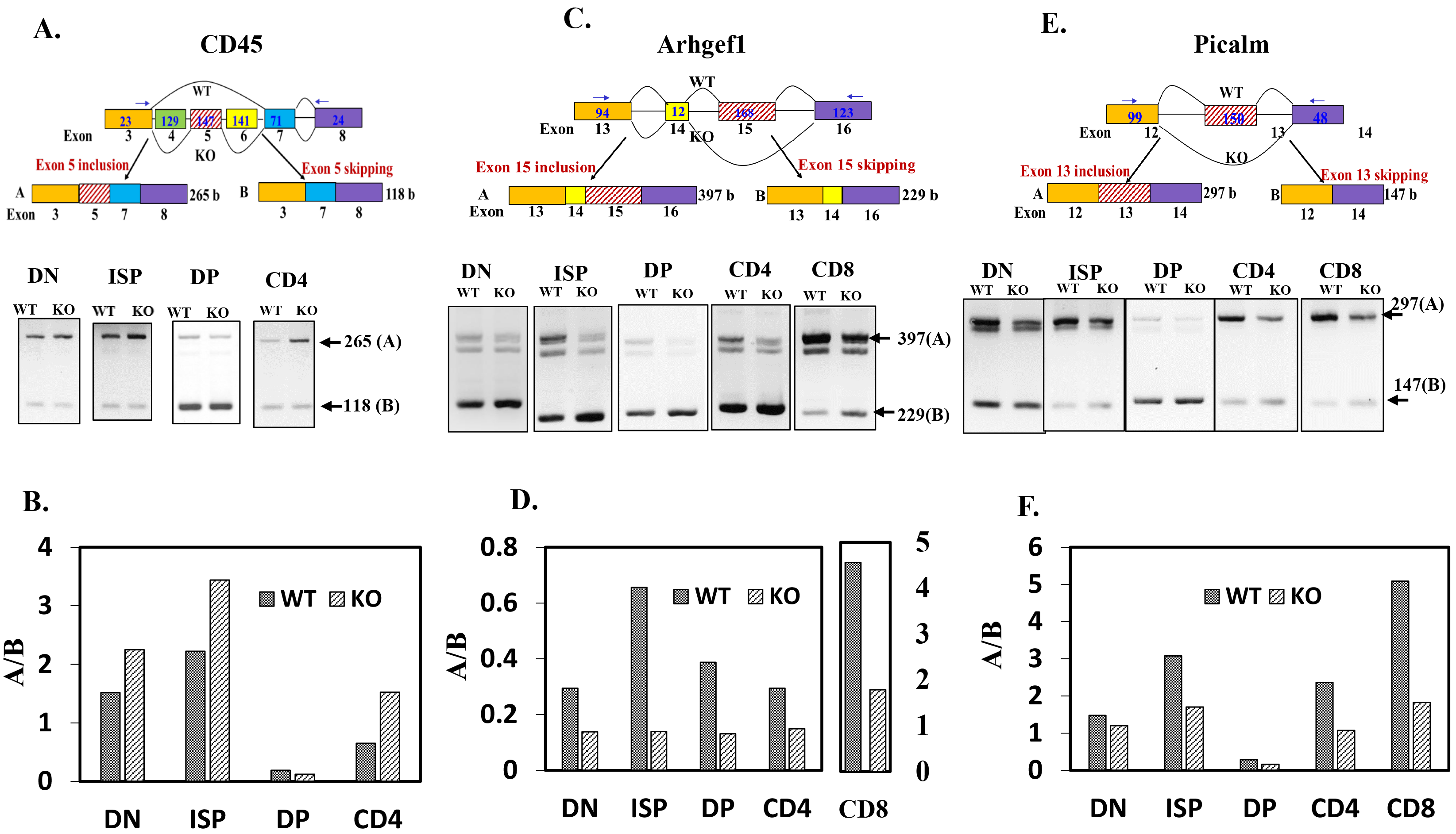
RT-PCR validation of RNA-seq data. RNA from the different thymocyte subpopulations was subjected to RT-PCR for the indicated genes. (A) and (B), CD45; (C) and (D), Arhgef1; (E) and (F), Picalm. (A),(C) and (E) Upper panels: schematic diagrams depicting partial gene structure of the alternatively spliced genes. Rectangular boxes represent the exons and the horizontal straight lines connecting the boxes represent the introns; the numbers below the boxes refer to the exon number of the gene and numbers inside the boxes refer to the length of the exons; the numbers within the terminal exons do not refer to the actual exon length but the length amplifiable by the RT-PCR primers; the arrow heads show the approximate binding positions of the RT-PCR primers; boxes with hashed lines show the alternative exons; curved lines connecting the boxes depict the splicing pattern. WT and KO refers to the splicing pattern prevalent in either the WT or the KO thymocytes as determined by RNA-seq analysis. Lower panels: ethidium bromide stained agarose gels showing RT-PCR products derived from total RNA from BRD4 WT and KO thymocytes. (B), (D) and (F) Histograms of the RT-PCR results; the ratios A/B (ratio of included exon transcript/ skipped exon transcript) were used as measure of alternative splicing.

Deletion of BRD4 changes the patterns of splicing of all three genes. For example, the frequency of inclusion of exon 5 of CD45 is increased in DN, ISP and CD4 cells in the absence of BRD4 (Fig.2A, B). In contrast, the frequencies of inclusion of exons 15 of Arhgef1 and exon 13 of Picalm are decreased in all subpopulations in the absence of BRD4 (Fig. 2C-F). Thus, BRD4 contributes to the regulation of splicing *in vivo* in all thymocyte subpopulations.

### Loss of BRD4 in T-ALL cells results in changes in splicing patterns

The growth of T cell acute lymphoblastic leukemia (T-ALL) is severely inhibited by either the inhibition of BRD4 binding to chromatin with a small molecule inhibitor, JQ1, or degron-mediated (dBET6) degradation (24). Accordingly, either treatment down-regulated a large number of transcripts (24). To determine whether BRD4 regulates splicing patterns in T-ALL, we analyzed the published T-ALL datasets (24). Analysis of the ChIP-seq data reveals that the effect of BRD4 degradation on BRD4 binding to chromatin is markedly greater than that of JQ1 treatment (Supplementary Figure 2). Since JQ1 functions by blocking BRD4 binding to chromatin through its bromodomains, these results suggest an association of BRD4 to chromatin independent of its bromodomains. In particular, JQ1 has relatively little effect on BRD4 binding within the gene body whereas dBET6 eliminates about 90% of the binding, consistent with the interpretation that BRD4 functions within the gene body independent of its binding to chromatin through its bromodomains.

Analysis of the T-ALL RNA-seq datasets (24) further revealed that treatment of T-ALL with either JQ1 or dBET6 resulted in large alterations in the splicing patterns (Figure 3). Consistent with its greater effect on RNA expression profiles and more severe growth inhibition of T-ALL cells, dBET6 treatment had a markedly larger effect on the number of splice events than did JQ1, although the overall distributions among the different splice forms were similar (Figure 3A, B). Skipped exon events represented over half of the changes resulting from either treatment; the distribution of skipped exon events was fairly even across the metagene (Supplementary Figure 3). Mutually exclusive exons represented the second largest category of change (Figure 3A, B). Alternative splicing events are enriched in genes that are differentially expressed in response to JQ1 or dBET6 (Figure 3C). Importantly, alternative splicing events are enriched among genes bound by BRD4 (Figure 3D). Analysis of the differentially spliced genes in both dBET6 and JQ1 treated cells shows that cell cycle pathways are preferentially affected (Supplementary Figure 4). Taken together, these results demonstrate that BRD4 contributes to the regulation of splicing.

**Figure 3.**
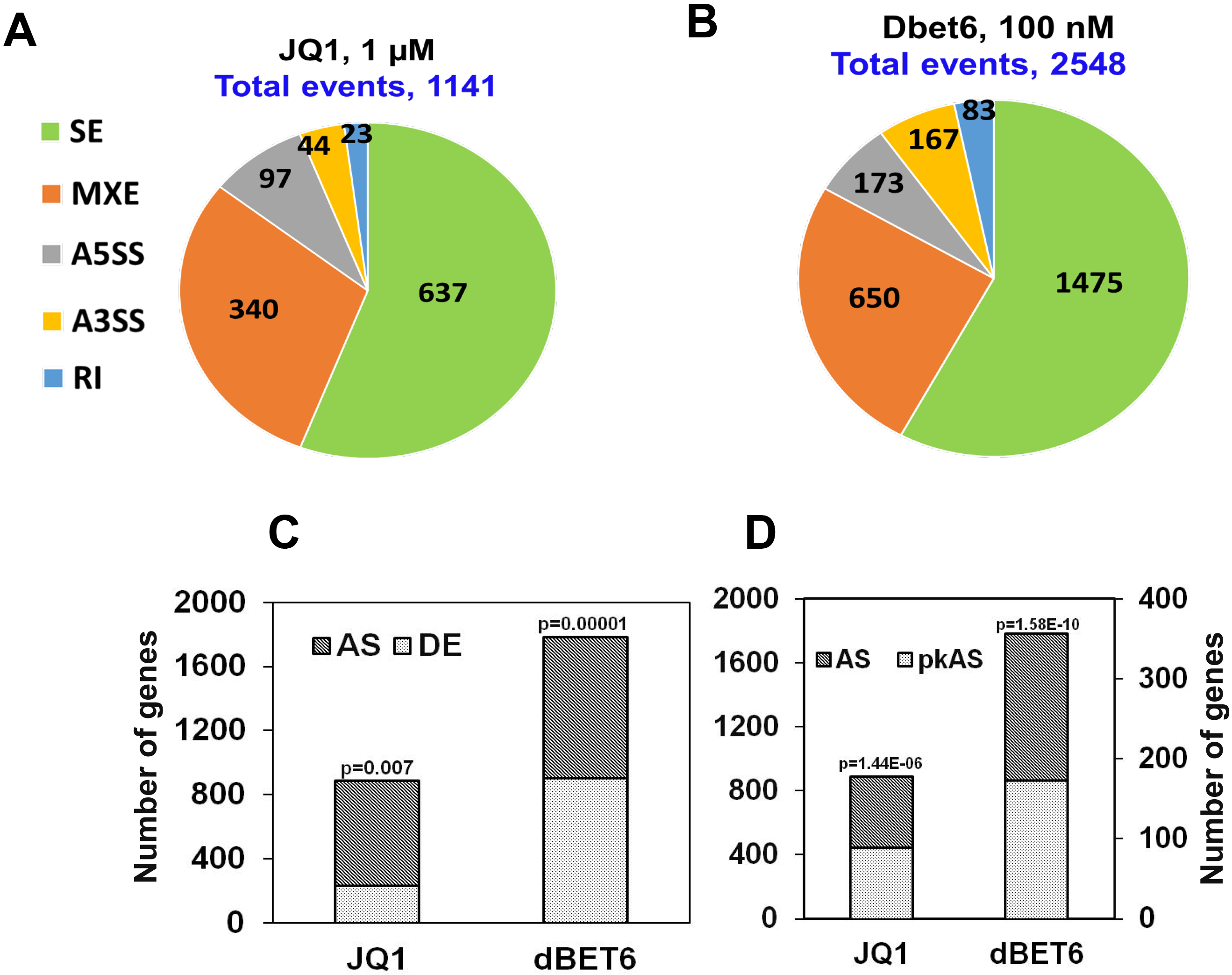
BET inhibition (JQ1) or BET degradation (dBET6) alter splicing patterns in T cell acute lymphoblastic leukemia (T-ALL) cells. (A) and (B) Pie chart showing distribution of different types of alternative splice events among the total of differentially spliced events in response to JQ1 treatment (A) and dBET6 treatment (B) in T-ALL cells. (C) Histogram showing proportion of alternative splice events (AS) among genes that are differentially expressed (DE) in response to JQ1 or dBET6 treatment. (D) Histogram showing proportion of alternative splice events (AS) among genes that are bound by BRD4 at the TSS (pkAS).p-values were obtained using a Hypergeometric test which tests the probability that the frequency AS genes derived from DE genes (overlap) is larger than expected from the population; a low p-value suggests the enrichment of AS genes in DE genes. Note for all the histograms, the number of genes showing changes in alternative splicing is plotted on the left-hand primary Y-axis with same scale while the number of genes with BRD4 peaks (D) is plotted on the secondary Y-axis on the right using a different scale. The total number of BRD4 peaks detected at TSS across the genome was 1123.

### BRD4 modulates splicing of the insulin receptor minigene

We examined the effect of over-expressing BRD4 on the splicing pattern of an insulin receptor (IR) minigene which has been extensively used in splicing studies (26). The minigene construct, inserted downstream of the CMV promoter, consists of exons 10-12, of which exon 11 is alternatively spliced (Fig 4A). Transfection of the IR minigene into HeLa cells results in the generation of two transcripts: a 310 bp RNA containing exons 10-11-12 and a 274 bp RNA containing only exons 10 and 12 (Fig. 4B). Co-transfection of the IR minigene with a BRD4 expression construct results in over-expression of BRD4 and an increase in the relative amount of exon 11 inclusion by about 1.5-fold (Fig. 4B,C, D). Thus, BRD4 expression correlates with altered patterns of alternative splicing.

**Figure 4.**
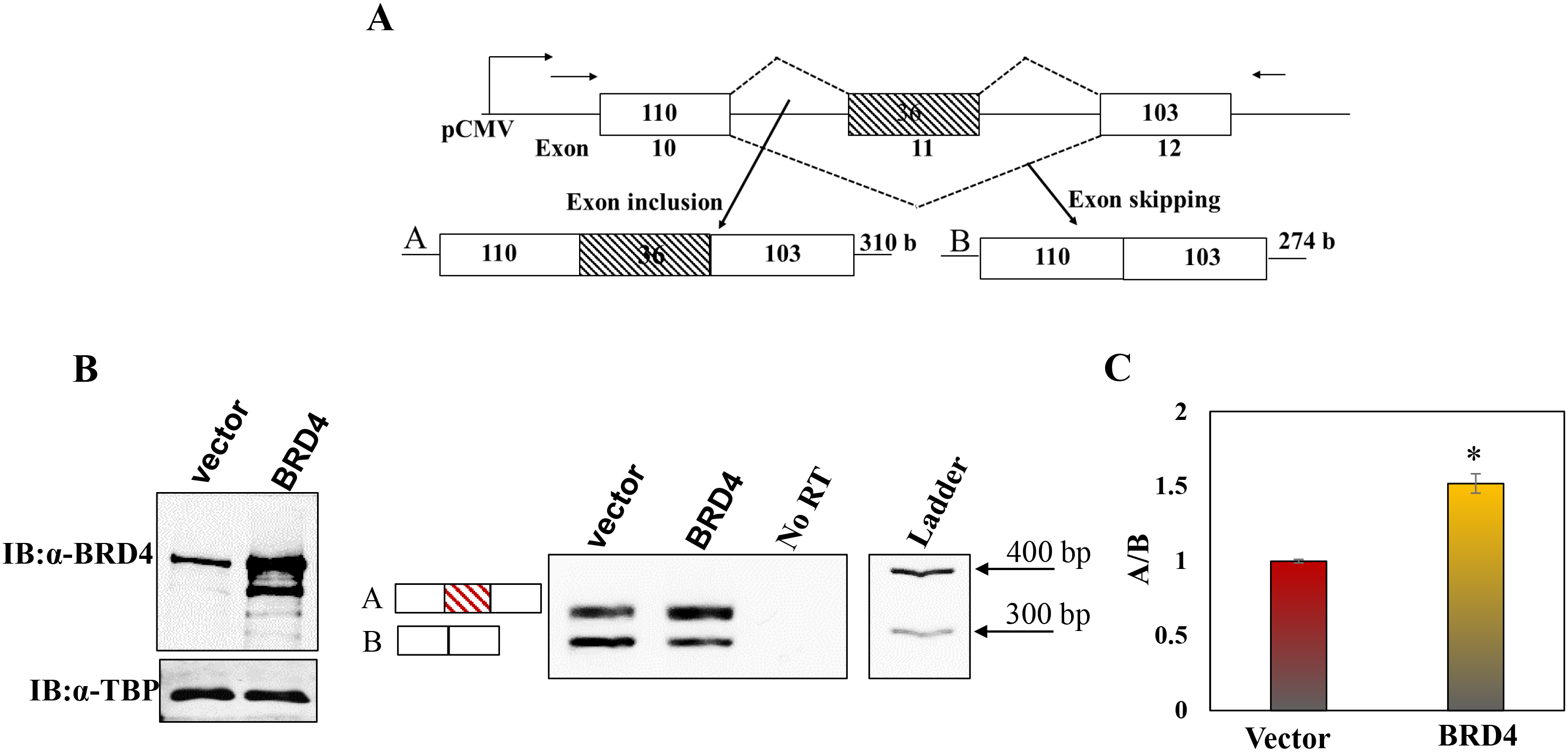
BRD4 regulates alternative splicing of the insulin receptor mini gene in Hela cells. (A) Schematic diagram of insulin receptor mini gene. The rectangular boxes represent exons and horizontal straight lines connecting the boxes represent the introns; the numbers below the boxes refer to the exon number of the gene and numbers inside the boxes refer to the length of the exons; the arrow heads show the approximate binding positions of the RT-PCR primers. The products of the two alternative exon usages are shown below. (B) BRD4 overexpression alters ratio of splice products. HeLa cells were cotransfected with the insulin receptor minigene and human WT BRD4 or control vector. Immunoblot of total HeLa cell extracts to determine the levels of BRD4 expression; TBP was used as a normalization control (Left); RT-PCR products derived from total RNA of transfected HeLa cells were electrophoresed on ethidium bromide stained agarose gel (Right). (C) Histogram of the RT-PCR results. The ratios A/B (ratio of included exon transcript/ skipped exon transcript) were used as measure of alternative splicing. *P<0.02.

BRD4 is known to have two enzymatic activities: a kinase activity that phosphorylates a variety of substrates, including Ser2 of the carboxy terminal domain (CTD) of Pol II and a histone acetyl transferase that acetylates the N-terminal tails of H3 and H4, as well as the core lysine, K122, of H3 (2, 3). Since both Ser2 phosphorylation of the CTD and histone modifications can affect splicing, we considered the possibility that the effect of BRD4 on splicing was indirectly through one of these mechanisms. To determine whether either of these enzymatic activities contributed to BRD4-mediated regulation of splicing, HeLa cells were co-transfected with the IR minigene and BRD4 mutants deficient in kinase or HAT activity (Supplementary Fig. 1). Transfection of the kinase mutant into HeLa cells resulted in a level of increased exon inclusion in the minigene indistinguishable from that of the WT BRD4 (Supplementary Fig. 1A-D). The finding that splicing is not affected by the absence of kinase activity indicates that BRD4 regulation is not mediated exclusively through its phosphorylation of the Pol II CTD. Similarly, expression of the BRD4 HAT mutant does not affect the pattern of splicing relative to the wild type BRD4, indicating that histone acetylation by BRD4 is not necessary for its effect on splicing (Supplementary Fig. 1E-G). Taken together, these results indicate that BRD4 regulation of splicing is a novel function, independent of either of its known enzymatic activities.

### BRD4 directly interacts with the splicing machinery

Although the above results demonstrate that BRD4 contributes to determining patterns of splicing, they do not distinguish whether that effect is indirect or direct through an interaction with the splicing machinery. Since previous mass spec analysis of factors that co-immunoprecipitate with BRD4 had identified a large series of splicing factors (21), we sought to validate and extend these analyses. To determine whether BRD4 interacts with splicing factors, HeLa cell nuclear extracts were subjected to immunoprecipitation with anti-BRD4 antibody and the ability to co-immunoprecipitate splicing factors assessed by immunoblotting. As shown in Fig. 5A, anti-BRD4 co-precipitates a series of splicing factors including components of the spliceosome, U1-70 and U1-A, as well as the regulatory splicing factors HNRNPM and Fus. Conversely, immunoprecipitation of HeLa extracts with anti-Fus antibody co-immunoprecipitated BRD4, as well as other spliceosome components, HNRNPM, U1-70 and U1-A (Fig.5B). Thus, BRD4 interacts with the splicing complex.

**Figure 5.**
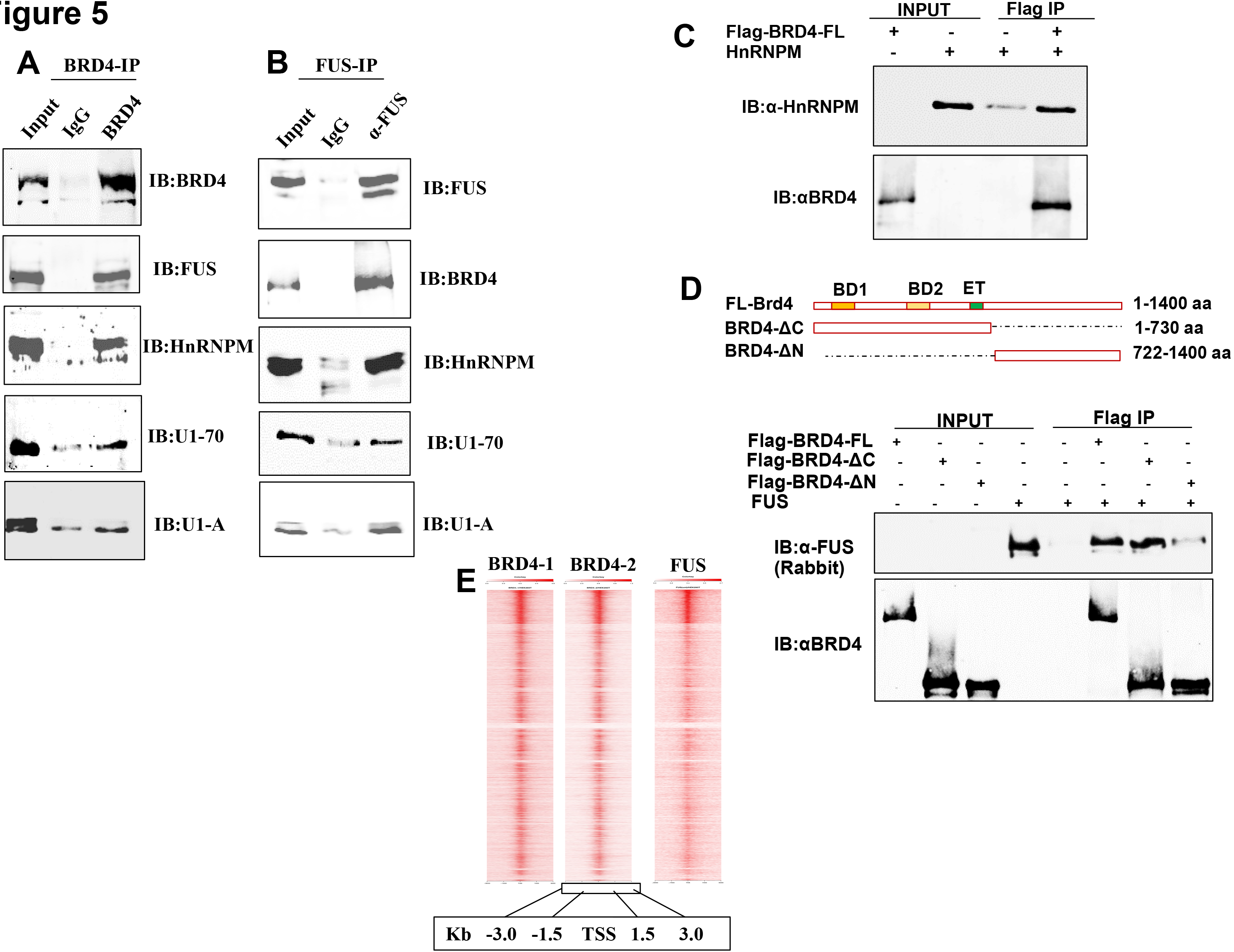
BRD4 interacts with splicing factors in vivo. (A) Immunoblot of BRD4-immunoprecipitate from HeLa nuclear extract with indicated antibodies to splicing factors Fus, HNRNPM, U1-70 and U1-A. (B) Immunoblot of Fus-immunoprecipitate from HeLa nuclear extract with indicated antibodies to BRD4 and splicing factors HNRNPM, U1-70 and U1-A. (C) Map showing Full length WT BRD4, Flag-tagged ΔC and ΔN BRD4 mutants. (D) Immunoblots showing pulldown analysis of BRD4-HNRNPM interaction. HNRNPM (0.25 μg) was pulled down with flag-BRD4 (0.5 μg) immobilized on Flag beads. (E) Immunoblots showing pulldown analysis of BRD4-FUS interaction. FUS (0.25 μg) was pulled down with flag-BRD4 (0.5 μg) or equimolar amounts of BRD4 mutants immobilized on Flag beads. (F) Enrichment Heat map showing co-localization of BRD4 with Fus across the genome.

Within the splicing complex, BRD4 binds directly to HnRNPM and Fus, as shown by pull-down experiments with recombinant proteins (Fig. 5C-D). Both rHnRNPM and rFus were efficiently pulled down by BRD4 (Fig. 5C-D). Using N-terminal and C-terminal truncations of BRD4, the binding site of Fus was mapped to the N-terminus (Fig. 5D). Taken together, these results are consistent with the interpretation that BRD4 contributes to the patterns of alternative splicing through its binding to components of the splicing machinery.

### BRD4 and Fus co-localize on the genome

The above results demonstrate that BRD4 interacts with splicing factors, but they do not indicate whether the interaction occurs on chromatin or in the nucleoplasm. Published ChIP-seq data sets of BRD4 (27) and Fus (10) were re-analyzed to assess the extent of co-localization on chromatin. The analysis revealed that BRD4 and Fus are highly co-localized on chromatin within 1.5 Kb of the transcription start site (TSS) and also across the gene body (Figure 5E, Supplementary Figure 6), consistent with a role in splicing.

### BRD4 co-localizes with splicing factors in situ

To directly visualize the association of BRD4 with the spliceosome *in situ,* HeLa cells were subjected to a proximity ligation assay (PLA) which detects proteins situated within 30-40nm of each other within the cell. As shown in Fig. 6, BRD4 is detected robustly in proximity to the splicing factors Fus, HnRNPM, U1-A and U1-70, but not with the negative control, nucleolin (Fig. S6). Taken together, these results document the colocalization of BRD4 with the spliceosome complex *in situ.*

**Figure 6.**
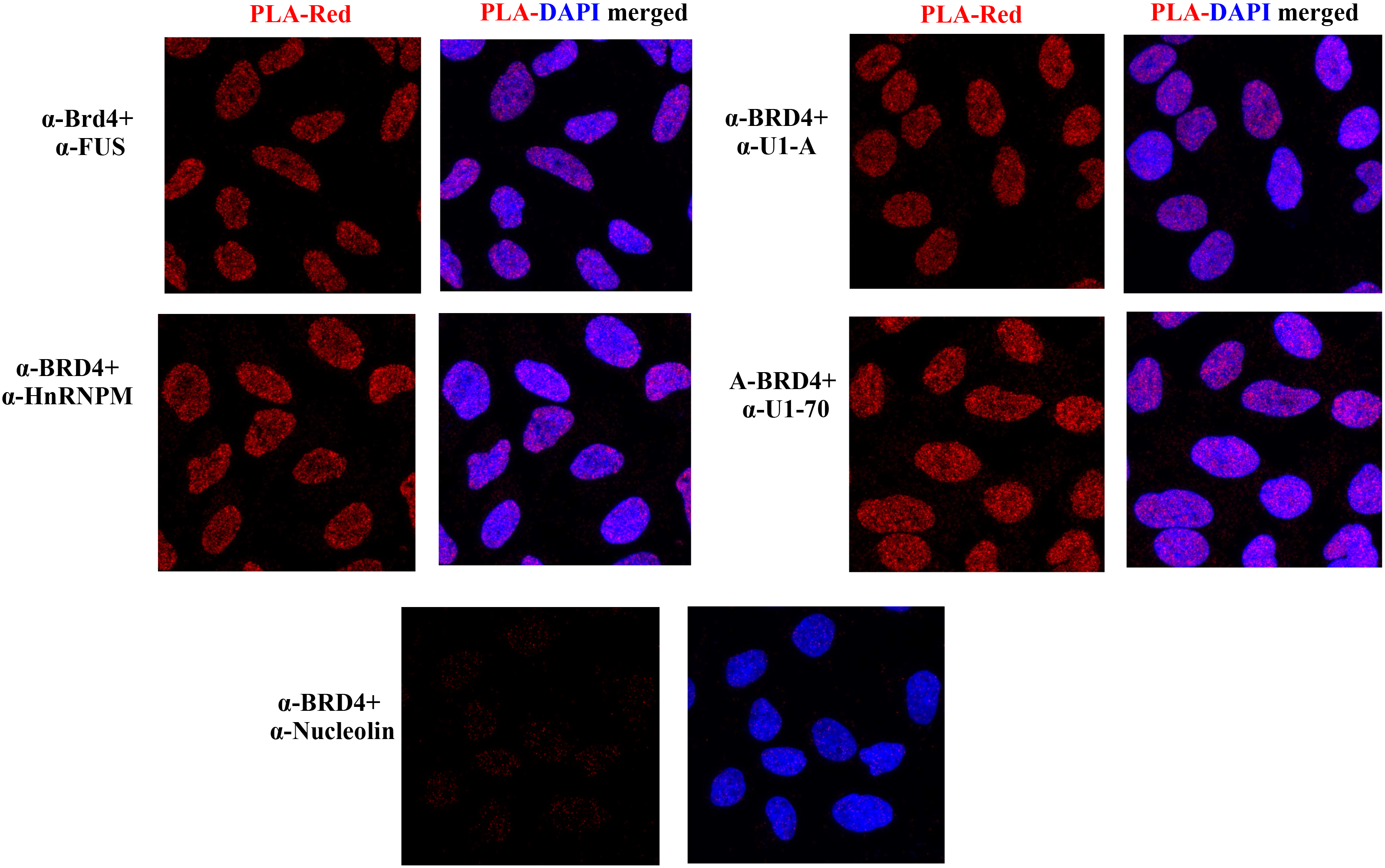
BRD4 interacts with other splicing factors *in situ.* Proximity ligation assays (PLA) were performed with anti-BRD4 and the antibodies for the indicated splicing factors on fixed HeLa cells. PLA with nucleolin was included as a negative control. PLA interaction is shown in red; DAPI staining in blue.

## Discussion

BRD4 is a member of the BET family of proteins that has been implicated in a variety of solid and hematological malignancies, as well as inflammatory diseases (1). Although BRD4 is being actively studied as a therapeutic target, knowledge of its biological functions is still incomplete. It is known that BRD4 serves as a scaffold for a variety of transcription factors, notably the transcription elongation factor PTEFb, delivering it to sites of transcription (28). Importantly, BRD4 plays an active role in regulating gene expression by linking chromatin structure and transcription through its HAT and kinase activities, respectively (2, 3). In the present study, we have established that BRD4 contributes to the regulation of splicing in both normal and cancer cells. The effect of BRD4 on splicing is likely to be direct since BRD4 is associated with the splicing machinery *in vivo* and co-localizes with it on the genome where it acts preferentially by modulating exon usage. Based on our findings, we hypothesize that BRD4 functions to integrate the processes of chromatin structure, transcription and splicing to ensure proper regulation of gene expression.

In the normal developing thymus, the stages of thymocyte differentiation - DN, ISP, DP and SP - are characterized by their cell surface expression of the markers CD4 and CD8. We have found that the transitions from one stage to another are accompanied by changes in the patterns of splicing, suggesting that splicing is developmentally regulated in the thymus. BRD4, which is expressed at approximately equal levels in each of these stages (25), contributes to this regulation since deletion of BRD4 results in changes in the splicing patterns, preferentially affecting exon skipping events, but not in expression of splicing factors (25).

Consistent with its role in cancer, BRD4 also contributes to patterns of alternative splicing in T-ALL cells, largely affecting the genes involved in cell cycle regulation. Interestingly, degradation of BRD4 results in much larger changes in splicing patterns than the inhibition of BRD4 binding to chromatin through its bromodomains. This suggests that BRD4 mediated regulation of alternative splicing does not depend entirely on its binding to chromatin through its bromodomains. Further support for this perspective comes from the observation that blocking of BRD4 bromodomain-mediated interaction with chromatin is largely confined to the promoter regions of genes whereas degradation of BRD4 targets both promoters and gene bodies. Whether it binds to chromatin at gene bodies directly by a bromodomain independent mechanism or indirectly through its binding to either the transcription elongation factors or splicing machinery remains to be determined.

In previous studies, we showed that the BRD4 HAT activity acetylates histones, particularly H3K122, resulting in nucleosome eviction and chromatin remodeling (2). BRD4 intrinsic kinase activity phosphorylates Ser2 of the Pol II CTD and is associated with transcriptional pause release (3). Since histone modifications and chromatin structure both have been implicated in regulating splicing, and phosphorylated Ser2 serves to recruit splicing factors (15, 29), these earlier studies suggested that BRD4 could regulate splicing indirectly. However, mutation of either of its enzymatic activities did not alter the pattern of splicing of the insulin receptor minigene, suggesting that the contribution of BRD4 to splicing is independent of these two activities. Although we cannot exclude the possibility that BRD4 contributes to splicing indirectly, our findings indicate that BRD4 regulation of splicing is independent of either of its enzymatic activities.

Mutations in splicing factors have been reported in a number of different malignancies, including myelodysplastic syndromes, acute myeloid leukemia, lung, breast and pancreatic cancers, pointing to a critical role of splicing in cancer (30–33). The splicing factor, HnRNPM, regulates the splicing pattern of CD44, which is known to play a role in cancer metastasis (34). BRD4 similarly plays a role in metastasis (35, 36). Translocations of the regulator of splicing, Fus, lead to sarcomas (37). More recently, an analysis of splicing patterns in nearly 9000 tumors revealed extensive alternative splicing associated with transformation (38). Surprisingly, in some tumors, the extent of novel splicing events was disproportionate to the mutational burden, suggesting that another regulatory mechanism may be at play (38). Based on our present finding that BRD4 directly interacts with both HnRNPM and Fus, it is tempting to speculate that BRD4, which is frequently overexpressed in cancer, exerts its effect on tumorigenesis and metastasis at least in part through its regulation of splicing. Future studies will be directed at addressing these possibilities.

## Methods

### Cell culture, Plasmids and Transient Transfections

HeLa cells were cultured in DMEM medium supplemented with 10% fetal bovine serum. A detailed list of the constructs used for transfections and preparation of recombinant proteins is provided in Supplemental Methods. The Insulin receptor mini gene plasmid was a gift from Dr Stefans Stamm (University of Kentucky) (26). Transient co-transfections of BRD4 constructs with the insulin receptor minigene plasmid were done in HeLa cells using Lipofectamine (Invitrogen), as described by the manufacturer. The transfected cells were harvested after 18 h and analyzed for protein levels and alternative splicing as described below.

### Alternative splicing detection using RT-PCR

For mini gene assays, reverse transcription (RT) was performed on purified total RNA using insulin receptor specific reverse primers, as described in Supplemental Methods. For alternative splicing validation of thymocytes RNA-seq data, 30 ng of total RNA from WT and KO thymocytes (25) was used with gene specific primers (0.1 μM). Primer sequences are shown in supplementary Table S3.

### Immunoprecipitation (IP) and Immunoblotting

For co-immunoprecipitation, 100 μg of HeLa nuclear extract was incubated with protein G agarose beads (Thermo Scientific) coated with 5 μg of appropriate antibody [anti-BRD4 antibody (A301-985A50, Bethyl laboratories) or anti-FUS antibody (Abcam)] overnight at 4 °C. To test for Flag-BRD4 direct binding to GST-FUS or HnRNPM, Flag beads (Anti-flag M2 Affinity Gel, Sigma) were incubated with equimolar concentrations of proteins for 3 hrs at 4 °C. Following incubation and washing, proteins were separated on SDS/PAGE gels and transferred onto a nitrocellulose membrane (GE Healthcare). For in vivo co-immunoprecipitation, the membrane was partitioned into three horizontal strips to probe multiple proteins with different molecular weight. After blocking the membrane with 5% fat-free milk in TBST, the blots were incubated with the primary antibodies BRD4 antibody, anti-FUS mouse monoclonal antibody, anti-FUS rabbit polyclonal antibody. HnRNP M1-4 mouse monoclonal IgG1 antibody, anti-U1A antibody, anti-U1-70K mouse monoclonal antibody, anti-flag rabbit polyclonal antibody, anti-TBP antibody. Catalog numbers and sources are provided in the Supplementary Methods. The secondary antibodies IRDye®800CM or IRDye®680RD (LI-COR Biosciences) were used for protein detection. All immunoblot analyses were performed using the Odyssey infrared scanner and secondary antibodies from Li-Cor.

### Proximity ligation assay (PLA)

For PLA, approximately 10^4^ Hela cells were grown overnight in μ-Slide Angiogenesis (Ibidi). PLA was conducted using the Duolink® In Situ PLA® Kit (Sigma) according to the manufacturer’s protocol. The primary antibodies used were as follows: anti-BRD4 rabbit polyclonal antibody, anti-FUS mouse monoclonal antibody, anti-FUS rabbit polyclonal antibody. HnRNP mouse monoclonal IgG1 antibody, anti-U1A antibody, anti-U1-70K Mouse monoclonal antibodyand anti-Nucleolin. Antibody dilutions, catalog numbers and sources are provided in the Supplementary Methods. Cells were observed with Zeiss LSM880 Multi-Photon Confocal Microscope.

### Bioinformatics analysis

Analysis of alternative splicing in thymocyte subpopulation was performed on the datasets reported in Gegonne et al. For BRD4-FUS co-localization analysis (Fig. 4), the following GEO datasets were used: BRD4 chip-seq samples BRD4-1, BRD4-2 in HEK293T cells using GEO dataset GSE51633 (27) and FUS chip-seq for Si-control, Si-FUS in HEK293T cells using GEO dataset PRJNA185008 (10). GEO dataset GSE79290 was used for alternative splicing and BRD4 ChIP-seq analysis in T-ALL cells (24). RNA-seq datasets corresponding to the 6 hr time point after JQ1 and dBET6 treatment in T-ALL cells were used for alternative splicing and differential expression analysis.

For RNA-seq analysis, sequencing reads were aligned to the reference genome (mouse: mm10 and human: hg19) with STAR aligner. Differential gene expression analysis was performed with DESeq2; a gene was considered as differentially expressed with its fold-change > 1.5 and FDR adjusted p-value < 0.01, as previously described (25). Alternative splicing was analyzed by rMATS (39); significant events were defined as events using junction counts with its FDR adjusted p-value < 0.05 and absolute value of inclusion level difference (IncLevelDifference) > 0.1. For ChIP-seq analysis, sequencing reads were aligned to the human reference genome hg19 by using Bowtie2. The duplicated reads were removed and only uniquely mapped reads were used for peak identification. ChIP-seq peaks were called using MACS2 (40) using default parameters. The enrichment analysis was performed using hypergeometric test.

### Statistical analysis

The data are presented as means ± standard deviation (S.D.) of at least three independent experiments. The p values indicate statistical significance, which is obtained using student’s t-test (two-tailed, unpaired).

## Acknowledgements

The authors gratefully acknowledge Drs. Ranjan Sen, Dan Larson, Ananda Roy and Richard Hodes for their critical review of the manuscript, the rest of the Singer lab for helpful discussions and Michael Kruhlak for assistance with imaging. This work was supported by the Intramural Research Program of the NIH, National Cancer Institute, Center for Cancer Research.

## REFERENCES

1. Hajmirza A, et al. (2018) BET Family Protein BRD4: An Emerging Actor in NFkappaB Signaling in Inflammation and Cancer. Biomedicines 6(1).

2. Devaiah BN, et al. (2016) BRD4 is a histone acetyltransferase that evicts nucleosomes from chromatin. Nature structural & molecular biology 23(6):540–548.

3. Devaiah BN, et al. (2012) BRD4 is an atypical kinase that phosphorylates serine2 of the RNA polymerase II carboxy-terminal domain. Proceedings of the National Academy of Sciences of the United States of America 109(18):6927–6932.

4. Baranello L, et al. (2016) RNA Polymerase II Regulates Topoisomerase 1 Activity to Favor Efficient Transcription. Cell 165(2):357–371.

5. Saldi T, Cortazar MA, Sheridan RM, & Bentley DL (2016) Coupling of RNA Polymerase II Transcription Elongation with Pre-mRNA Splicing. Journal of molecular biology 428(12):2623–2635.

6. Will CL & Luhrmann R (2011) Spliceosome structure and function. Cold Spring Harbor perspectives in biology 3(7).

7. Brugiolo M, Herzel L, & Neugebauer KM (2013) Counting on co-transcriptional splicing. F1000prime reports 5:9.

8. Oesterreich FC, et al. (2016) Splicing of Nascent RNA Coincides with Intron Exit from RNA Polymerase II. Cell 165(2):372–381.

9. Kornblihtt AR, de la Mata M, Fededa JP, Munoz MJ, &Nogues G (2004) Multiple links between transcription and splicing. RNA (New York, N.Y.) 10(10):1489–1498.

10. Schwartz JC, et al. (2012) FUS binds the CTD of RNA polymerase II and regulates its phosphorylation at Ser2. Genes & development 26(24):2690–2695.

11. Huang Y, Yao X, & Wang G (2015) ‘Mediator-ing’ messenger RNA processing. Wiley interdisciplinary reviews. RNA 6(2):257–269.

12. Wei WJ, et al. (2012) YB-1 binds to CAUC motifs and stimulates exon inclusion by enhancing the recruitment of U2AF to weak polypyrimidine tracts. Nucleic acids research 40(17):8622–8636.

13. Ruiz-Velasco M, et al. (2017) CTCF-Mediated Chromatin Loops between Promoter and Gene Body Regulate Alternative Splicing across Individuals. Cell systems 5(6):628–637.e626.

14. Guillouf C, Gallais I, & Moreau-Gachelin F (2006) Spi-1/PU.1 oncoprotein affects splicing decisions in a promoter binding-dependent manner. The Journal of biological chemistry 281(28): 19145–19155.

15. Naftelberg S, Schor IE, Ast G, & Kornblihtt AR (2015) Regulation of alternative splicing through coupling with transcription and chromatin structure. Annual review of biochemistry 84:165–198.

16. Venkataramanan S, Douglass S, Galivanche AR, & Johnson TL (2017) The chromatin remodeling complex Swi/Snf regulates splicing of meiotic transcripts in Saccharomyces cerevisiae. Nucleic acids research 45(13):7708–7721.

17. Chen W, Luo L, & Zhang L (2010) The organization of nucleosomes around splice sites. Nucleic acids research 38(9):2788–2798.

18. Zhou HL, et al. (2011) Hu proteins regulate alternative splicing by inducing localized histone hyperacetylation in an RNA-dependent manner. Proceedings of the National Academy of Sciences of the United States of America 108(36):E627–635.

19. Fiszbein A & Kornblihtt AR (2016) Histone methylation, alternative splicing and neuronal differentiation. Neurogenesis (Austin, Tex.) 3(1):e1204844.

20. Berkovits BD, Wang L, Guarnieri P, & Wolgemuth DJ (2012) The testis-specific double bromodomain-containing protein BRDT forms a complex with multiple spliceosome components and is required for mRNA splicing and 3′-UTR truncation in round spermatids. Nucleic acids research 40(15):7162–7175.

21. Rahman S, et al. (2011) The Brd4 extraterminal domain confers transcription activation independent of pTEFb by recruiting multiple proteins, including NSD3. Molecular and cellular biology 31(13):2641–2652.

22. Hussong M, et al. (2017) The bromodomain protein BRD4 regulates splicing during heat shock. Nucleic acids research 45(1):382–394.

23. Kanno T, et al. (2014) BRD4 assists elongation of both coding and enhancer RNAs by interacting with acetylated histones. Nature structural & molecular biology 21(12):1047–1057.

24. Winter GE, et al. (2017) BET Bromodomain Proteins Function as Master Transcription Elongation Factors Independent of CDK9 Recruitment. Molecular cell 67(1):5–18.e19.

25. Gegonne A, et al. (2018) Immature CD8 Single-Positive Thymocytes Are a Molecularly Distinct Subpopulation, Selectively Dependent on BRD4 for Their Differentiation. Cell reports 24 (1):117–129.

26. Kosaki A, Nelson J, & Webster NJ (1998) Identification of intron and exon sequences involved in alternative splicing of insulin receptor pre-mRNA. The Journal of biological chemistry 273(17):10331–10337.

27. Liu W, et al. (2013) Brd4 and JMJD6-associated anti-pause enhancers in regulation of transcriptional pause release. Cell 155(7):1581–1595.

28. Patel MC, et al. (2013) BRD4 coordinates recruitment of pause release factor P-TEFb and the pausing complex NELF/DSIF to regulate transcription elongation of interferon-stimulated genes. Molecular and cellular biology 33(12):2497–2507.

29. Luco RF, Allo M, Schor IE, Kornblihtt AR, & Misteli T (2011) Epigenetics in alternative pre-mRNA splicing. Cell 144(1):16–26.

30. Qiu J, et al. (2016) Distinct splicing signatures affect converged pathways in myelodysplastic syndrome patients carrying mutations in different splicing regulators. RNA (New York, N.Y.) 22(10):1535–1549.

31. de Necochea-Campion R, Shouse GP, Zhou Q, Mirshahidi S, & Chen CS (2016) Aberrant splicing and drug resistance in AML. Journal of hematology & oncology 9(1):85.

32. Martinez-Montiel N, Anaya-Ruiz M, Perez-Santos M, & Martinez-Contreras RD (2017) Alternative Splicing in Breast Cancer and the Potential Development of Therapeutic Tools. Genes 8(10).

33. Wang BD, et al. (2017) Alternative splicing promotes tumour aggressiveness and drug resistance in African American prostate cancer. Nature communications 8:15921.

34. Xu Y, et al. (2014) Cell type-restricted activity of hnRNPM promotes breast cancer metastasis via regulating alternative splicing. Genes & development 28(11):1191–1203.

35. Hu Y, et al. (2015) BRD4 inhibitor inhibits colorectal cancer growth and metastasis. International journal of molecular sciences 16(1):1928–1948.

36. Alsarraj J & Hunter KW (2012) Bromodomain-Containing Protein 4: A Dynamic Regulator of Breast Cancer Metastasis through Modulation of the Extracellular Matrix. International journal of breast cancer 2012:670632.

37. Crozat A, Aman P, Mandahl N, & Ron D (1993) Fusion of CHOP to a novel RNA-binding protein in human myxoid liposarcoma. Nature 363(6430):640–644.

38. Kahles A, et al. (2018) Comprehensive Analysis of Alternative Splicing Across Tumors from 8,705 Patients. Cancer cell 34(2):211–224.e216.

39. Shen S, et al. (2014) rMATS: robust and flexible detection of differential alternative splicing from replicate RNA-Seq data. Proceedings of the National Academy of Sciences of the United States of America 111(51):E5593–5601.

40. Zhang Y, et al. (2008) Model-based analysis of ChIP-Seq (MACS). Genome biology 9(9):R137.

